# Sclerostin directly stimulates osteocyte synthesis of fibroblast growth factor-23

**DOI:** 10.1101/2020.10.29.360024

**Authors:** Nobuaki Ito, Matthew Prideaux, Asiri R. Wijenayaka, Dongqing Yang, Renee T. Ormsby, Lynda F. Bonewald, Gerald J. Atkins

## Abstract

Osteocyte produced fibroblast growth factor 23 (FGF23) is the key regulator of serum phosphate (Pi) homeostasis. The interplay between parathyroid hormone (PTH), FGF23 and other proteins that regulate FGF23 production and serum Pi levels is complex and incompletely characterised. Evidence suggests that the protein product of the *SOST* gene, sclerostin (SCL), also a PTH target and also produced by osteocytes, plays a role in FGF23 expression, however the mechanism for this effect is unclear. Part of the problem of understanding the interplay of these mediators is the complex multi-organ system that achieves Pi homeostasis *in vivo*. In the current study, we sought to address this using a unique cell line model of the osteocyte, IDG-SW3, known to express FGF23 at both the mRNA and protein levels. In cultures of differentiated IDG-SW3 cells, both PTH_1-34_ and recombinant human (rh) SCL remarkably induced *Fgf23* mRNA expression dose-dependently within 3 hours. Both rhPTH_1-34_ and rhSCL also strongly induced C-terminal FGF23 protein secretion. Secreted intact FGF23 levels remained unchanged, consistent with constitutive post-translational cleavage of FGF23 in this cell model. Both rhPTH_1-34_ and rhSCL treatments significantly suppressed mRNA levels of *Phex, Dmp1* and *Enpp1* mRNA, encoding putative negative regulators of FGF23 levels, and induced *Galnt3* mRNA expression, encoding N-acetylgalactosaminyl-transferase 3 (GalNAc-T3), which protects FGF23 from furin-like proprotein convertase-mediated cleavage. The effect of both rhPTH_1-34_ and rhSCL was antagonised by pre-treatment with the NF-κβ signalling inhibitors, BAY11 and TPCK. RhSCL also stimulated *FGF23* mRNA expression in *ex vivo* cultures of human bone. These findings provide evidence for the direct regulation of FGF23 expression by sclerostin. Locally expressed sclerostin via the induction of FGF23 in osteocytes thus has the potential to contribute to the regulation of Pi homeostasis.

## Introduction

Fibroblast growth factor family member 23 (FGF23) is a protein hormone produced by osteoblasts and osteocytes, and is a potent regulator of blood levels of Pi [1, 2]. The expression of both FGF receptor 1 (FGFR1) and αKlotho forms a receptor complex allowing responsiveness to FGF23 in organs remote from its synthesis, most notably the renal proximal tubule epithelium [3–5]. Both Pi and the active hormonal form of vitamin D, 1α, 25-dihydroxyvitamin D3 (1,25D) have been shown to stimulate the synthesis of FGF23 *in vivo* and *in vitro* [6–12], and FGFR1 expression by osteocytes has been shown to play a pivotal role in sensing the serum Pi level [13]. Furthermore, other gene products are clearly implicated in regulating the synthesis of FGF23, since mutations of these genes leads to its dysregulation. Specifically, mutations in the genes phosphate regulating endopeptidase homolog, X-linked (*PHEX*), dentin matrix protein 1 (*DMP1*) and ectonucleotide pyrophosphatase/phosphodiesterase I (*ENPP1*) result in hypophosphatemic rickets with inappropriately high FGF23 levels, in both humans and genetically modified mice, however the mechanism by which they do so is unknown.

In addition to regulating serum calcium levels, parathyroid hormone (PTH) is also known to suppress serum Pi levels *via* the down regulation of the sodium phosphate co-transporter-2a (NaPi-2a) in renal proximal tubules: this molecule is also a target of FGF23. PTH also stimulates the synthesis of 1,25D by increasing the expression of *CYP27B1* and decreasing the expression of *CYP24A1* [14, 15]. In contrast, FGF23 down-regulates 1,25D by decreasing *CYP27B1* and increasing *CYP24A1* [16]. PTH and FGF23 are therefore now regarded as the dominant regulators of serum 1,25D. In addition to the interaction of FGF23 and PTH in regulating serum Pi and calcium levels, there is accumulating evidence that they regulate each other’s production. Lavi-Moshayoff and co-workers [17] showed that parathyroidectomy prevented the increase of FGF23 with high Pi levels seen in the adenine high-phosphorus diet-induced early kidney failure mouse model, which supports the hypothesis that PTH can directly stimulate FGF23 production. In the same report, it was shown that PTH increased *Fgf23* mRNA levels in the rat osteoblastic cell line, UMR-106. Subsequently, Rhee and colleagues [18] reported that mice, in which a specific constitutively active PTH receptor 1 (PTHR1) was over-expressed specifically in osteocytes (Dmp1-caPTHR1), displayed high serum levels of intact FGF23, together with high bone levels of *Fgf23* mRNA and of the gene *Galnt3*. The significance of the latter finding is that the protein product of *Galnt3*, N-acetylgalactosaminyl-transferase 3 (GalNAc-T3), is critical in the FGF23 regulatory pathway, as it protects intact FGF23 protein from inactivation by cleavage, by initiating *O*-glycosylation adjacent to the cleavage site [19–21]. These authors also showed that PTH increased levels of *Fgf23* mRNA in primary osteoblast/osteocyte cultures [18]. In addition, they showed that PTH or PTHR1 signalling down-regulated *Sost* mRNA expression in rat UMR-106 cells, in the bone of Dmp1-caPTHR1 mice, and in primary osteoblasts and osteocytes, consistent with the findings of others [22, 23]. The effects of PTH or PTHR1 signalling on FGF23 expression were antagonised by co-treatment with sclerostin, the product of the *Sost* gene, in UMR-106 cells or by co-overexpression of *Sost* in osteocytes (Dmp1-SOST tg) in Dmp1-caPTHR1 mice [17, 18]. In humans, *SOST* is the causative gene for high bone mass condition Sclerosteosis (#269500) [24]. Sclerostin was shown to block canonical Wnt signalling by binding to the Wnt co-receptors low density lipoprotein receptor-related proteins (LRP)-5 and −6 [25]. Therefore, it was hypothesised that FGF23 production is directly stimulated by canonical Wnt signalling and indirectly by PTH *via* its suppressive effect on *SOST* gene expression. Ren and colleagues [26] treated hypophosphatemic *Dmp1* knockout mice with neutralising antibody to SCL, which improved osteomalacia in these animals without attenuating the hypophosphatemia or the high levels of circulating FGF23. However, Ryan *et al*. [27] reported that serum intact FGF23 levels in *Sost* gene knock-out mice were reduced, consistent with a positive rather than a negative effect of sclerostin on FGF23 expression. Also, we have reported that *Sost* mRNA levels are elevated in the long bones of *Phex* deletion (Hyp) mice [28], which show hypophosphatemic rickets with increased serum intact FGF23 when compared with normal controls, although others have reported decreased levels in neonatal animals [29].

To date, *in vitro* studies to investigating the regulation of FGF23 have been conducted in UMR-106 or primary bone cells, as the only cells expressing detectable FGF23 *in vitro* [11, 17, 30]. Recently, the mouse osteoblast-like cell line, IDG-SW3, was established, which differentiates into osteocyte-like cells expressing FGF23 [12, 31, 32]. To further investigate the regulation of FGF23 by PTH and sclerostin, we used differentiated IDG-SW3 cells, in which PTH and sclerostin both induced robust up-regulation of *Fgf23* mRNA expression.

## Materials and Methods

### IDG-SW3 cells

IDG-SW3 cells were cultured as described previously [12, 31, 32] and differentiated into cells that were morphologically osteocyte-like and expressed an osteocytic gene profile over a 35-day period. All further experiments were performed on IDG-SW3 cells differentiated for 35 days.

### Recombinant PTH and Sclerostin treatment

Differentiated IDG-SW3 cells were treated for the indicated time intervals with rhPTH_1-34_ (Tocris Bioscience) at concentrations between 1-100 nM or recombinant human sclerostin (rhSCL) (R&D Systems) at concentrations between 1-50 ng/ml [28, 33] for the time periods indicated. Media were also collected from differentiated IDG-SW3 cells treated for 24h with rhPTH_1-34_ (100 nM) or rhSCL (50 ng/ml) for FGF23-specific ELISA. To determine the effects of rhPTH_1-34_ and rhSCL on *Fgf23* or other related mRNA expression, differentiated IDG-SW3 cells treated with rhPTH_1-34_ (100 nM) or rhSCL (50 ng/ml) were harvested at 3, 6, 12 and 24 hours. In some experiments, cells were also treated with the NF-κß signalling inhibitor, BAY11 (Sigma) at 100 μM or N-Tosyl-L-Phenylalanine Chloromethyl Ketone (TPCK) (Sigma) at 100 μM one hour before the addition of rhPTH_1-34_ (100 nM) or rhSCL (50 ng/ml) and harvested after 6h.

### Human bone samples

Human cancellous bone samples were obtained with written consent from patients undergoing total hip replacement surgery. Bone samples were digested twice with collagenase (10 mg/ml) at 37°C for 30 min to remove marrow and lining cells [34], and then cut into pieces (~1 mm in diameter) using a sterile scalpel before being placed evenly in 12-well tissue culture plates in αMEM medium (Sigma) containing 2% FCS [32].

### RNA purification and quantitative real-time PCR (qPCR)

RNA was extracted from IDG-SW3 cells and human bone samples with TRIzol (Invitrogen) and then cDNA was synthesised with iScript (Bio-Rad). Real-Time PCR (CFX Connect™; Bio-Rad) was used to quantitate mRNA levels by qPCR, as described previously [35]. cDNA specific primers for qPCR were designed to amplify mRNA for murine glyceraldehyde-3-phosphate dehydrogenase (*Gapdh*), *Fgf23, Sost, Phex, Dmp1, Enpp1, Galnt3, IL-6* and human *GAPDH, FGF23, PHEX, DMP1* and *ENPP1*. All qPCR measurements were normalised to the expression of *Gapdh/GAPDH* mRNA (Supplementary Table S1).

### FGF23 ELISA

FGF23 protein levels in the media and cell lysates were analysed by two different types of ELISA: the FGF-23 (C-Term) ELISA Kit (Immutopics, San Diego, CA, USA) utilises two epitopes, both detecting the C-terminal side of the cleavage point such that this kit detects both full-length FGF23 and its C-terminal fragment. On the other hand, the Intact FGF-23 ELISA Kit (Kainos, Tokyo, Japan) has epitopes N and C terminal to the physiological cleavage point (amino acid 179-180), and therefore exclusively detects functional, full-length FGF23 [36].

### Statistical analysis

All experiments were performed in triplicate. All data shown represent the mean ± standard error of the mean (SEM). Statistical analyses were performed by using One-Way Analysis of Variance (ANOVA) followed by Tukey’s post-hoc test or Student’s t test (two-tailed). Data were analysed using GraphPad Prism software (GraphPad Prism, CA, USA). The significance level is indicated by asterisks: \**p* < 0.05; \*\**p* < 0.01; *** *p* < 0.001.

## Results

### Effect of PTH_1-34_ and sclerostin on FGF23 expression and secretion

Differentiated IDG-SW3 cells expressed abundant *Fgf23* mRNA, as reported previously [31]. When treated with rhPTH_1-34_ and rhSCL, *Fgf23* mRNA levels increased markedly, in a dosedependent manner. *Fgf23* mRNA levels increased by more than 700-fold in response to rhPTH_1-34_ (100 nM) and by more than 200-fold in response to rhSCL (50 ng/ml), compared to untreated controls, in 24 hours (Fig. 1A, B). Time-course studies revealed that *Fgf23* mRNA levels peaked at more than 2,000 and 3,000-fold for rhPTH 100 nM and rhSCL 50 ng/ml, respectively, within 3 to 12 hours of treatment (Fig. 1C, D). PTH and rhSCL also both induced *FGF23* mRNA expression in human cancellous bone samples cultured *ex vivo*, although to a lesser extent due possibly to the inefficient diffusion of these agents into the lacunocanalicular pores by passive diffusion (Suppl. Fig. S1).

**Figure 1:**
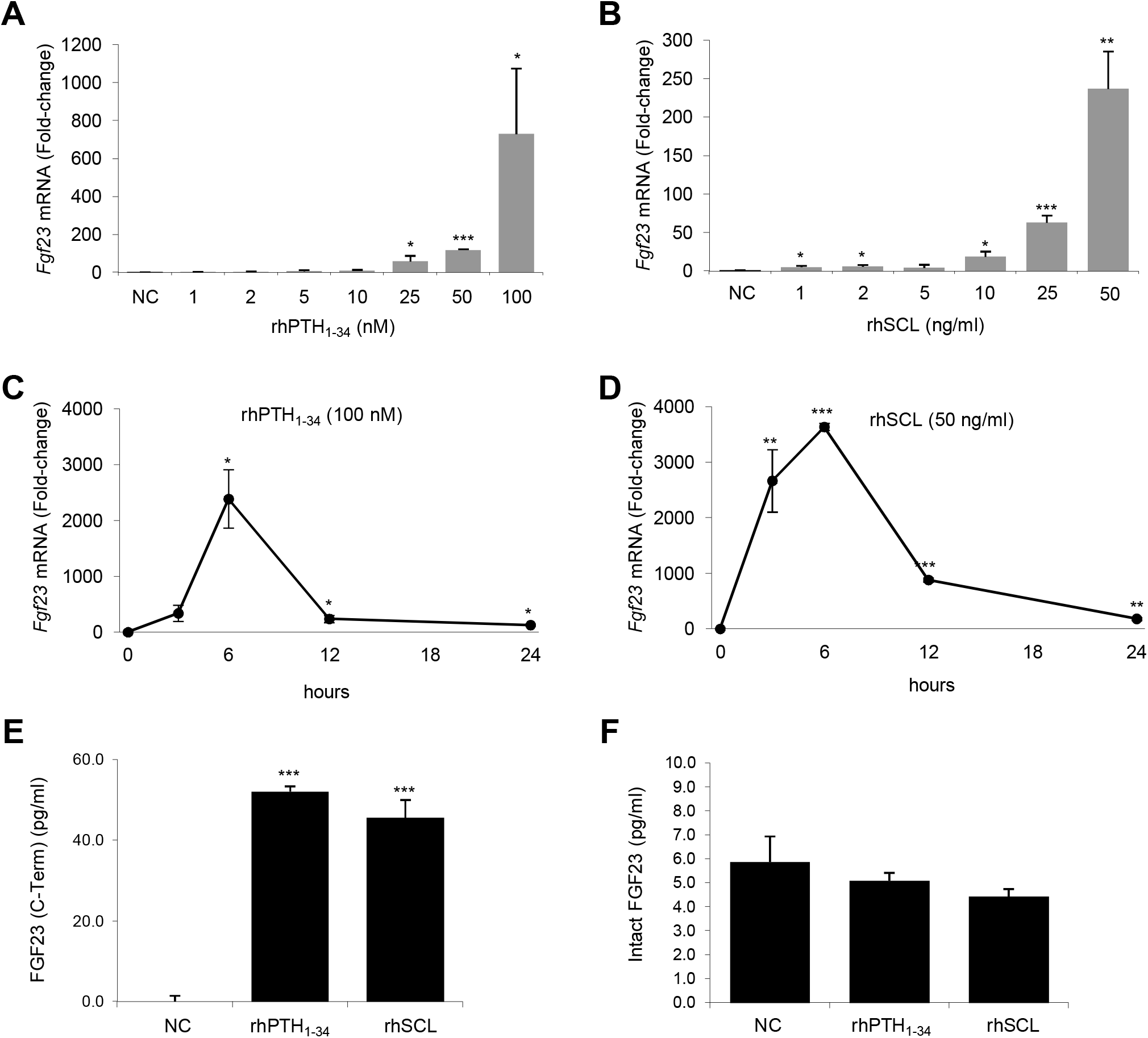
Effect of PTH and rhSCL treatment on *Fgf23* mRNA expression and FGF23 secretion. IDG-SW3 cells differentiated for 5 weeks were treated with A) rhPTH_1-34_ and B) rhSCL for 24 h at the concentrations indicated and *Fgf23* mRNA levels measured by real time RT-PCR. *Fgf23* mRNA levels were also measured in a time course in response to C) rhPTH_1-34_ (100 nM), and D) rhSCL (50 ng/ml). Gene expression was normalised to *Gapdh* mRNA levels and data described as mean fold-change ± standard error of the mean (SEM) relative to the untreated normal control (NC) or the time zero level. Media from IDG-SW3 cells differentiated for 5 weeks and treated with rhPTH_1-34_ (100 ng/ml) or rhSCL (50 ng/ml) for 24h were measured using E) FGF23 (C-Term) and F) Intact FGF23 ELISA kits. Data shown are means ± SEM of quadruplicate readings. NC: normal control; Significant difference to the reference level is depicted by \**p* < 0.05; \*\**p* < 0.01; \*\*\**p* < 0.001.

Treatment with either rhPTH_1-34_ or rhSCL for 24 hours resulted in abundant secretion of FGF23 measured by the C-terminal FGF23 ELISA kit (Fig. 1E) confirming that the effects on *Fgf23* mRNA levels by both PTH and rhSCL resulted in new FGF23 protein synthesis. However, there was no significant change in the levels of secreted intact FGF23 in response to these treatments (Fig. 1F). This is consistent with our previous observations, that the IDG-SW3 model exhibits constitutively active furin-like proprotein convertase activity and/or a deficiency in the ability to protect newly translated FGF23 protein by *O*-glycosylation [32].

### Effect of PTH and SCL on FGF23 regulatory gene expression

The expression of genes implicated in control of FGF23 levels was also investigated. In differentiated IDG-SW3 cells, *Phex* (Fig. 2A & B), *Dmp1* and *Enpp1* mRNA levels were significantly decreased by both rhPTH_1-34_ and rhSCL at several concentrations (Suppl. Fig. S2, S3), although *Dmp1* mRNA levels were up-regulated at lower concentrations of rhPTH_1-34_ (1 nM) and rhSCL (1 nM) (Suppl. Fig. S2A&B). Suppression of *Phex, Dmp1* and *Enpp1* mRNA by rhPTH_1-34_ and rhSCL was seen by 3 to 6 hours of treatment and persisted for at least 24 hours for *Phex* and *Dmp1* (Fig. 2C & D, Suppl. Fig. S2 C&D, S3 C&D). A clear although variable increase of *Galnt3* mRNA levels was observed in response to either rhPTH_1-34_ or rhSCL by 3-6 hours of treatment (Fig. 3A-D).

**Figure 2:**
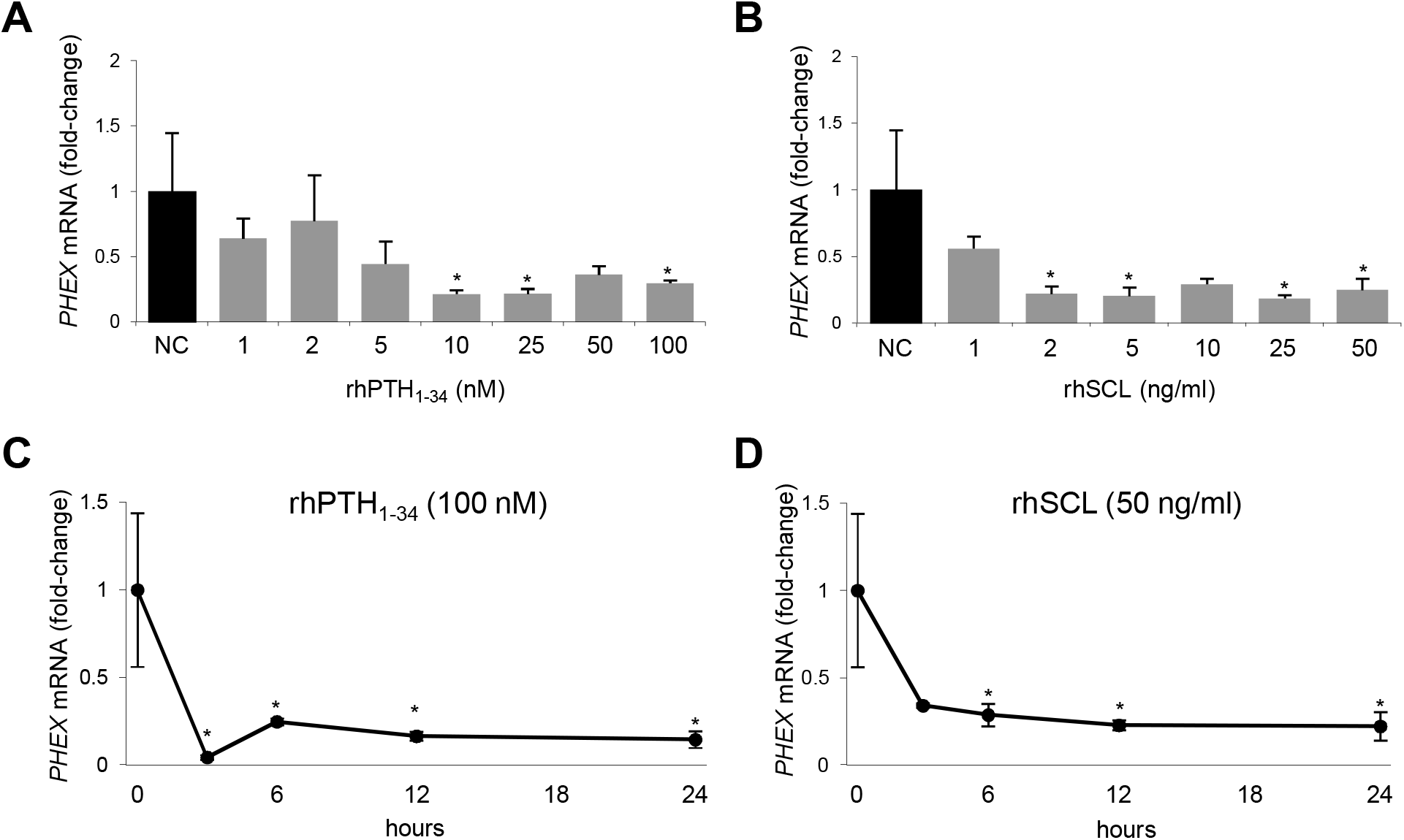
Effect of PTH and rhSCL treatment on *Phex* mRNA expression. IDG-SW3 cells differentiated for 5 weeks were treated with A) rhPTH_1-34_ and B) rhSCL for 24 h at the concentrations indicated and *Phex* mRNA levels measured by real time RT-PCR. *Phex* mRNA levels were also measured in a time course in response to C) rhPTH_1-34_ (100 nM), and D) rhSCL (50 ng/ml). Gene expression was normalised to *Gapdh* mRNA levels and data described as mean fold-change ± SEM relative to the untreated normal control (NC) or the time zero level. Significant difference to the reference level is depicted by \**p* < 0.05; \*\**p* < 0.01; \*\*\**p* < 0.001.

**Figure 3:**
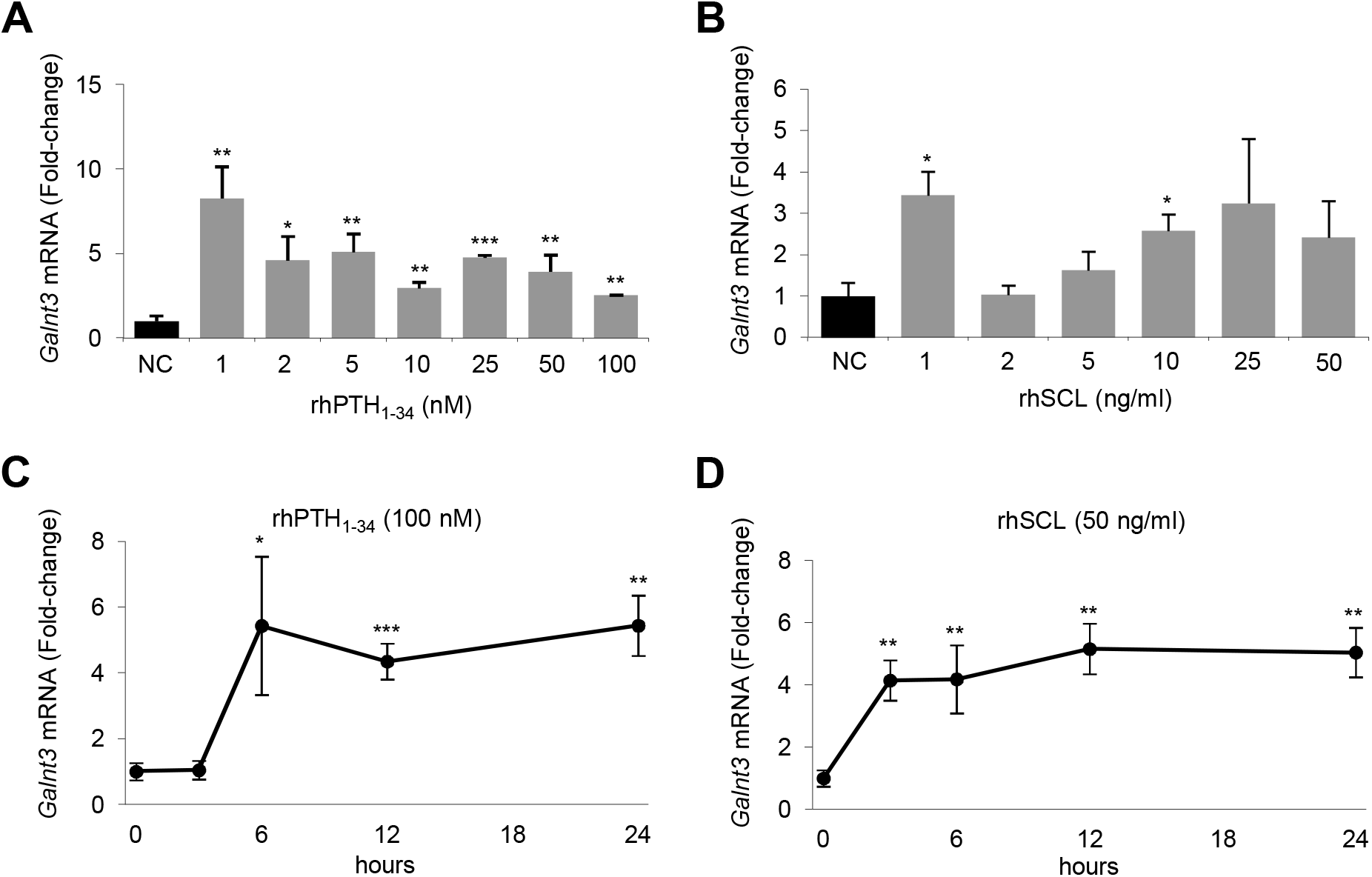
Effect of PTH and rhSCL treatment on *Galnt3* mRNA expression. IDG-SW3 cells differentiated for 5 weeks were treated with A) rhPTH_1-34_ and B) rhSCL for 24 h at the concentrations indicated and *Galnt3* mRNA levels measured by real time RT-PCR. *Galnt3* mRNA levels were also measured in a time course in response to C) rhPTH_1-34_ (100 nM), and D) rhSCL (50 ng/ml). Gene expression was normalised to *Gapdh* mRNA levels and data described as mean fold-change ± SEM relative to the untreated normal control (NC) or the time zero level. Significant difference to the reference level is depicted by \**p* < 0.05; \*\**p* < 0.01; \*\*\**p* < 0.001.

### Effect of NFκB inhibition

We reported recently that pro-inflammatory factors, including tumour necrosis factor alpha (TNFα) and interleukin-1β (IL-1β) stimulated the synthesis of *Fgf23* mRNA via NF-κß signalling [32], and it was also reported by others that PTH stimulates NF-κß signalling in bone cells and in renal tubular cells [37–39]. In the current study, the induction of *IL-6* mRNA, a well-known readout of NF-κß signalling, was observed in response to rhPTH_1-34_ and rhSCL (Fig. 4A-D). To investigate the involvement of NF-κß signalling in PTH and sclerostin mediated regulation on *Fgf23* expression, differentiated IDG-SW3 cells were treated with the NF-κß inhibitors, BAY11 or TPCK, one hour prior to rhPTH_1-34_ or rhSCL treatments. Both BAY11 and TPCK significantly antagonised the effect of rhPTH_1-34_ and rhSCL on *Fgf23* mRNA expression (Fig. 4E & F).

**Figure 4:**
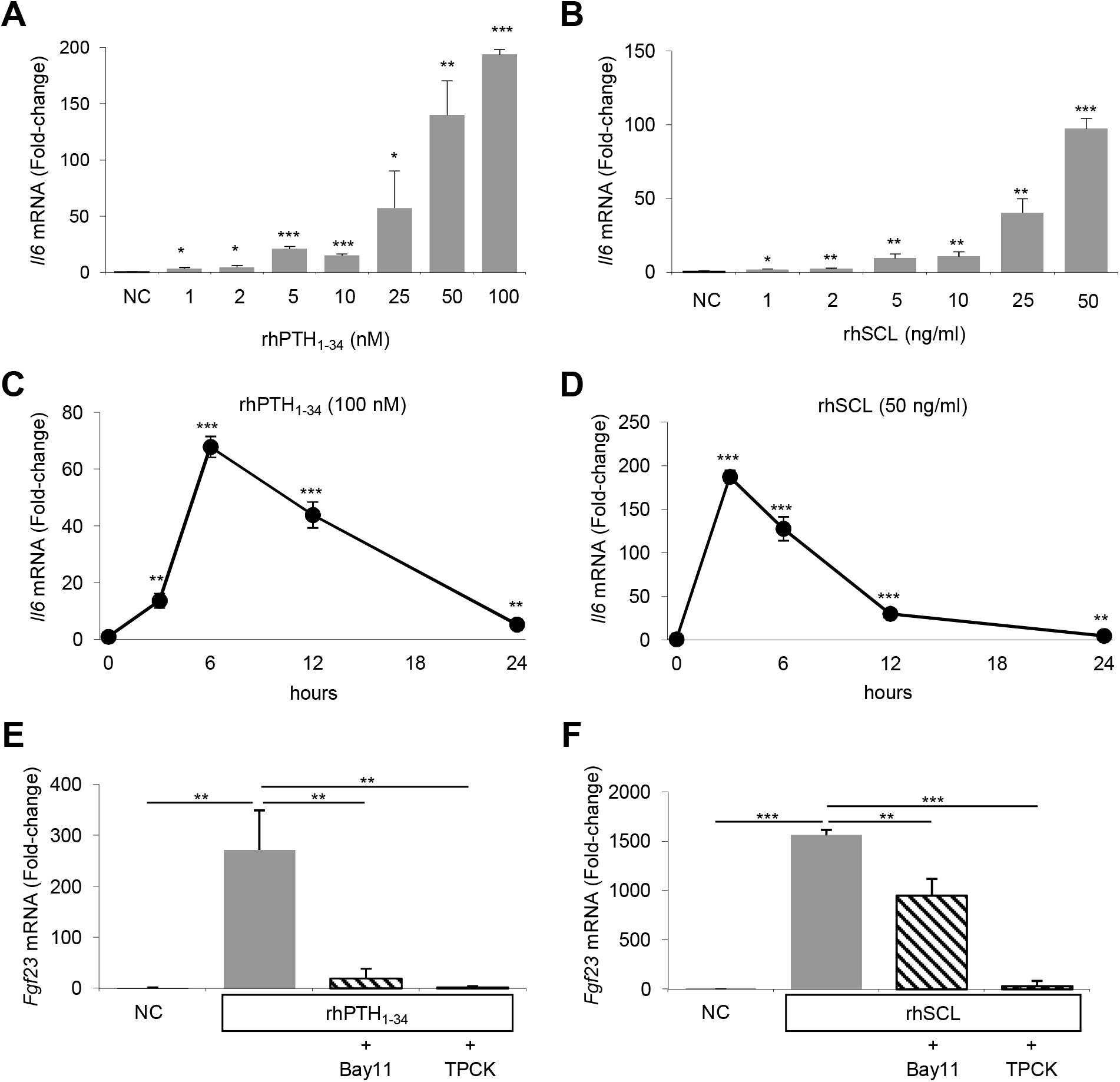
Effect of PTH and rhSCL on the IL-6/NF-κB pathway. IDG-SW3 cells differentiated for 5 weeks were treated with A) rhPTH_1-34_ and B) rhSCL for 24 h at the concentrations indicated and *Il6* mRNA levels measured by real time RT-PCR. *Il6* mRNA levels were also measured in a time course in response to C) rhPTH_1-34_ (100 nM), and D) rhSCL (50 ng/ml). Differentiated (5 wk) IDG-SW3 cells were pre-treated with NF-κβ signal inhibitors Bay11 (100 μM) or TPCK (100 μM) for 1h before treatment with E) rhPTH_1-34_ (100 nM) or F) rhSCL (50 ng/ml) for 24h and *Fgf23* mRNA levels were quantified by real-time RT-PCR. Gene expression was normalised to *Gapdh* mRNA levels and data described as mean fold-change ± SEM relative to the untreated normal control (NC) or the time zero level. Significant difference to the reference level is depicted by \**p* < 0.05; \*\**p* < 0.01; \*\*\**p* < 0.001.

## Discussion

PTH has been a long time candidate regulator of FGF23 levels. It has been shown that injection of rhPTH_1-34_ increased serum intact FGF23 levels within 18 hours [40] with no significant change in serum Pi levels. However, in that study, the effect on FGF23 levels was preceded by an increase in 1,25D levels, and, as stated, 1,25D is a known positive regulator of FGF23. Brown *et al*. [41] reported high intact FGF23 levels in a patient with Jansen’s metaphyseal chondrodysplasia (#156400), a condition caused by a gain of function mutation of the *PTH1R* gene resulting in the constitutive activation of the PTH and PTH related protein (PTHrP) signalling pathway. As this patient also showed low serum Pi and normal serum 1,25D levels, this finding implied a direct stimulatory effect of PTH on FGF23 expression [41]. These observations illustrate the complexity of unravelling the *in vivo* regulation of Pi, Ca and 1,25D levels, and whether PTH stimulates FGF23 directly or indirectly. Rhee *et al*. and Lavi-Moshayoff *et al*., respectively, showed the direct positive effect of PTH on *Fgf23* mRNA production possibly *via* the down-regulation of *Sost* expression in mouse primary osteoblast/osteocyte cultures and the rat osteoblast-like cell line UMR-106, which was previously the only cell line known to express *Fgf23* [17, 18, 30]. *SOST* is considered to be one of the key target genes when PTH1-34 is administered in an intermittent fashion as a pro-anabolic treatment for osteoporosis. We recently showed that sclerostin targets cells at the late osteoblast/pre-osteocyte and osteocyte stages, target populations to those of PTH, and acts as an autocrine/paracrine regulator of matrix mineralisation [28, 35, 42] and of osteocyte support of osteoclast activity [43]. It was possible, therefore, that the actions of PTH and sclerostin were overlapping, bisynchronous or conflicting, with respect to FGF23 production.

The murine osteoblast-like cell-line, IDG-SW3, has emerged as a promising cell model, with which to test the regulation of FGF23 expression, in particular when it is differentiated to a mature osteocyte-like stage [12, 31]. Here, *Fgf23* mRNA expression was strongly stimulated by PTH in these cells, up to more than 2,000-fold 6 hours after treatment. Importantly, synthesis of *Fgf23* mRNA was also induced by rhSCL, to a similar extent to rhPTH_1-34_. Under the experimental conditions employed, the *Sost* mRNA level was unchanged by rhPTH_1-34_ exposure (data not shown), which is inconsistent with previous reports using the MLO-A5 or UMR-106 cell lines, or mouse primary osteoblast/osteocytes [17, 18, 22, 23].

PHEX, DMP1 and ENPP1 proteins are thought to constitutively suppress serum intact FGF23 levels in healthy subjects [5]. Here, the observed down-regulation of expression of *Phex, Dmp1* and *Enpp1* mRNA and the increase of *Galnt3* mRNA levels in response to rhPTH_1-34_ or rhSCL, could all constitute part of the mechanism for the increase of *Fgf23* mRNA and/or FGF23 protein levels in response to rhPTH_1-34_, or indeed sclerostin *in vivo*, given the results of the loss of function mutation of these genes [20, 21, 36, 44–49].

As we have found previously that pro-inflammatory mediators increased *Fgf23* transcription *via* an NF-κβ-dependent pathway [32] and a downstream target gene of NF-κβ, *IL6*, was strongly induced by both rhPTH_1-34_ and rhSCL, NF-κβ signalling pathway inhibitors were used to determine if this pathway is also involved in the effect of PTH and sclerostin on *Fgf23* mRNA transcription. Two inhibitors, Bay11 and TPCK, significantly reversed the effect of both rhPTH_1-34_ and rhSCL on *Fgf23* mRNA levels. However, further studies will be required to fully elucidate the signalling pathways, by which these mediators regulate FGF23 expression.

The dramatic increase in *Fgf23* mRNA levels observed in response to both PTH_1-34_ and rhSCL, was reflected in the amount of secreted C-terminal fragments of FGF23 protein in the media. However, similar to our previous findings with pro-inflammatory mediators [32], intact FGF23 secreted in the media were unchanged, confirming that the levels of the intact hormone are finally determined by the level of protective post-translational *O*-glycosylation and Furin/Furin-like protein convertase activity, which mediates cleavage or protection of FGF23 protein [20, 21, 50–54]. This post-translational regulatory mechanism also functions as the terminal rate-limiting step to define the amount of active FGF23 secreted in response to serum Pi levels *in vivo* [51, 52, 54]. Consistent with the importance of this mechanism and with findings here, PTH_1-34_ was shown to elicit C-terminal FGF23 *in vivo* in mice without altering intact hormone levels, unless a furin inhibitor was also administered [55]. Although the rhPTH_1-34_ and rhSCL failed to increase intact FGF23 levels in IDG-SW3 cultures despite increased *Galnt3* mRNA expression, IDG-SW3 might be a promising tool to investigate the detailed mechanism of the Pi sensing system which remains to be fully elucidated beyond the identification of the pivotal role of FGFR1 expression by osteocytes in the process [13]. We have reported previously that at least *Fgf23* mRNA levels are sensitive to Pi using this cell model [12].

The findings of this study beg the question as to their possible physiological relevance. We previously reported that NF-κβ signalling in response to pro-inflammatory cytokines or LPS also induce the transcription of *Fgf23* mRNA without changing the intact FGF23 level in IDG-SW3 cells [32]. Therefore, it might be that the activators of osteoclast differentiation and activity, such as the local hypoxia due to low serum Fe [56] or NF-κβ signalling, also enhance the transcription of *FGF23* mRNA for the purpose of increasing intact FGF23 levels to decrease excess serum Pi due to bone resorption. We have demonstrated that sclerostin induces mineral release by osteocytes [35, 42], suggesting that local induction of FGF23 production may also occur in response to osteocytic osteolysis. Consistent with this, evidence for increased osteocytic osteolysis was reported in the murine XLH model, *Hyp* [57]. On the other hand, a continuously high serum level of PTH also has a bone catalytic effect [58] and sclerostin is identified as the inhibitor of osteogenic differentiation [24, 25, 28]. In light of this evidence, an osteoclastic or anti-osteogenic signal might have the ability to induce *FGF23* transcription with a concomitant suppressive effect on *PHEX, DMP1* and *ENPP1* mRNA and a stimulating effect on *GALNT3* expression, partly *via* the NF-κβ signalling pathway.

There are no reports of human sclerosteosis patients [24, 25] showing disturbance of serum Pi levels. However, *Sost* KO mice were reported to exhibit low serum intact FGF23 levels with high serum Pi [27]. Also, intriguingly, in Jansen’s metaphyseal chondrodysplasia patients caused by constitutively active PTH1R, both serum intact FGF23 and C-term FGF23 were reported to be high, which causes low serum Pi levels [41], and which may imply that sclerostin and PTH are not only inducers of *FGF23* mRNA transcription, as observed here, but also involved in the post-translational regulation of FGF23 *in vivo* to some extent. Consistent with this hypothesis is the observation that in patients with chronic hypoparathyroidism, hyperphosphatemia is not fully corrected by FGF23 action [59]. Interestingly, elevated serum sclerostin levels were reported in a cohort of young adult patients with XLH; unfortunately, FGF23 levels were not measured preventing a direct correlation between the two [60]. It will be of interest to observe the effects that clinical neutralisation of FGF23 in the case of XLH and other hypophosphatemic patient cohorts [61] has on either circulating or bone sclerostin levels.

In summary, here we report the potent induction of FGF23 transcription and translation by PTH and sclerostin. The precise post-translational regulatory mechanism(s) of intact FGF23 levels requires further study in terms of elucidating the Pi sensing mechanisms in osteocytes. To this end, IDG-SW3 is a promising cell model with which to study this. This study also provides the intriguing possibility that the down-regulation of *SOST* by PTH is required for anabolism in response intermittent PTH_1-34_ treatment because sclerostin and PTH have both overlapping and possibly discordant effects.

## Acknowledgments

This work was supported by funding from the National Health and Medical Research Council of Australia (NHMRC) Project Grant Scheme (Grant ID 10477960).

---

**Supplementary Figure S1:** Effect of PTH and rhSCL treatment on gene expression in human bone. Cancellous bone fragments prepared as described in Materials and Methods were treated with rhPTH_1-34_ (100 ng/ml) or rhSCL (50 ng/ml) for 24h. Bone was processed by total RNA and gene expression measured by real-time RT-PCR for A) *FGF23*, B) *PHEX*, C) *DMP1*, D) *ENPP1*, E) *GALNT3* mRNA expression. Gene expression was normalised to *GAPDH* mRNA levels and data described as mean fold-change ± SEM relative to the untreated normal control (NC). Significant difference to the reference level is depicted by * p<0.05; ** p<0.01; *** p<0.001.

**Supplementary Figure S2:** Effect of PTH and rhSCL treatment on *Dmp1* mRNA expression. IDG-SW3 cells differentiated for 5 weeks were treated with A) rhPTH_1-34_ and B) rhSCL for 24 h at the concentrations indicated and *Dmp1* mRNA levels measured by real time RT-PCR. *Dmp1* mRNA levels were also measured in a time course in response to C) rhPTH_1-34_ (100 nM), and D) rhSCL (50 ng/ml). Gene expression was normalised to *Gapdh* mRNA levels and data described as mean fold-change ± SEM relative to the untreated normal control (NC) or the time zero level. Significant difference to the reference level is depicted by * p<0.05; ** p<0.01; *** p<0.001.

**Supplementary Figure S3:** Effect of PTH and rhSCL treatment on *Enpp1* mRNA expression. IDG-SW3 cells differentiated for 5 weeks were treated with A) rhPTH_1-34_ and B) rhSCL for 24 h at the concentrations indicated and *Enpp1* mRNA levels measured by real time RT-PCR. *Enpp1* mRNA levels were also measured in a time course in response to C) rhPTH_1-34_ (100 nM), and D) rhSCL (50 ng/ml). Gene expression was normalised to *Gapdh* mRNA levels and data described as mean fold-change ± SEM relative to the untreated normal control (NC) or the time zero level. Significant difference to the reference level is depicted by * p<0.05; ** p<0.01; *** p<0.001.

**Supplementary Table S1:**
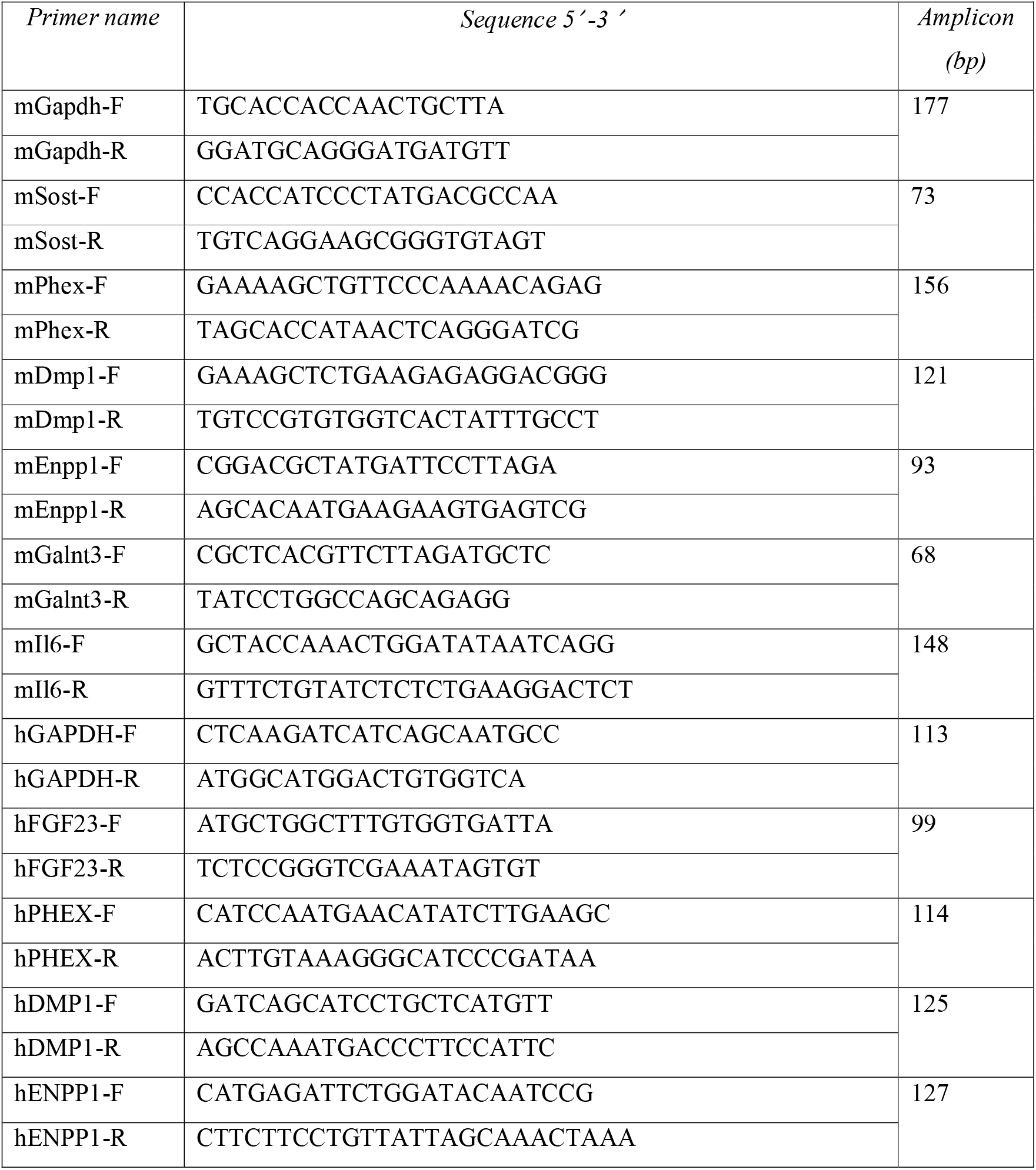
Real-time RT-PCR oligonucleotide primer sequences. At least one of each primer pair was designed to flank an intron-exon boundary and amplification confirmed to be mRNA-specific. Species is denoted by m (mouse) or h (human) followed by the gene name and direction of amplification F (forward) or R (reverse).

