## Supplementary Figure 1 for "Sclerostin directly stimulates osteocyte synthesis of fibroblast growth factor-23"

### Slide 1
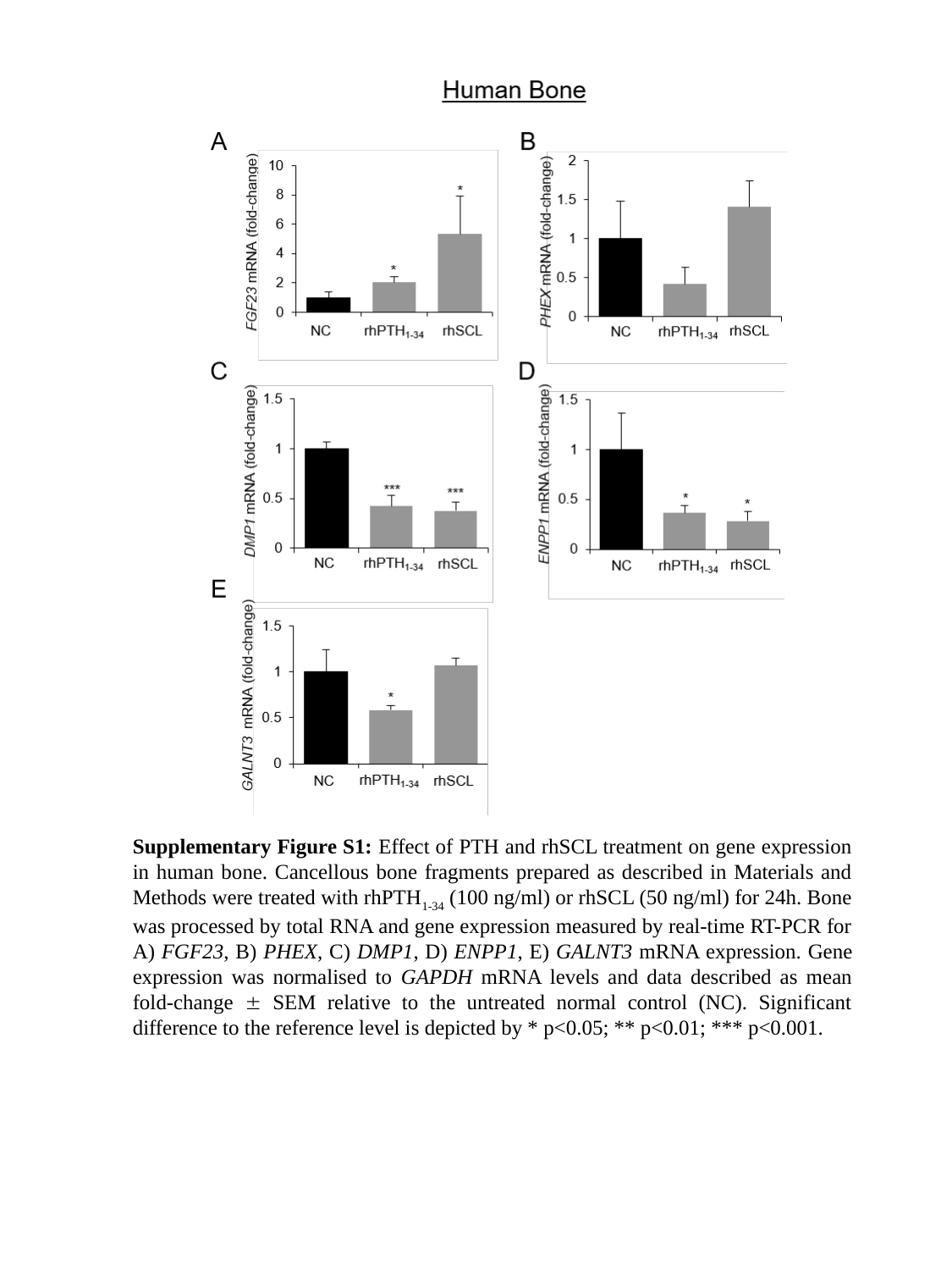

Supplementary Figure S1: Effect of PTH and rhSCL treatment on gene expression in human bone. Cancellous bone fragments prepared as described in Materials and Methods were treated with rhPTH_1-34_ (100 ng/ml) or rhSCL (50 ng/ml) for 24h. Bone was processed by total RNA and gene expression measured by real-time RT-PCR for A) FGF23, B) PHEX, C) DMP1, D) ENPP1, E) GALNT3 mRNA expression. Gene expression was normalised to GAPDH mRNA levels and data described as mean fold-change ± SEM relative to the untreated normal control (NC). Significant difference to the reference level is depicted by * p<0.05; ** p<0.01; *** p<0.001.
