## Supplementary Figure 2 for "Sclerostin directly stimulates osteocyte synthesis of fibroblast growth factor-23"

### Slide 1
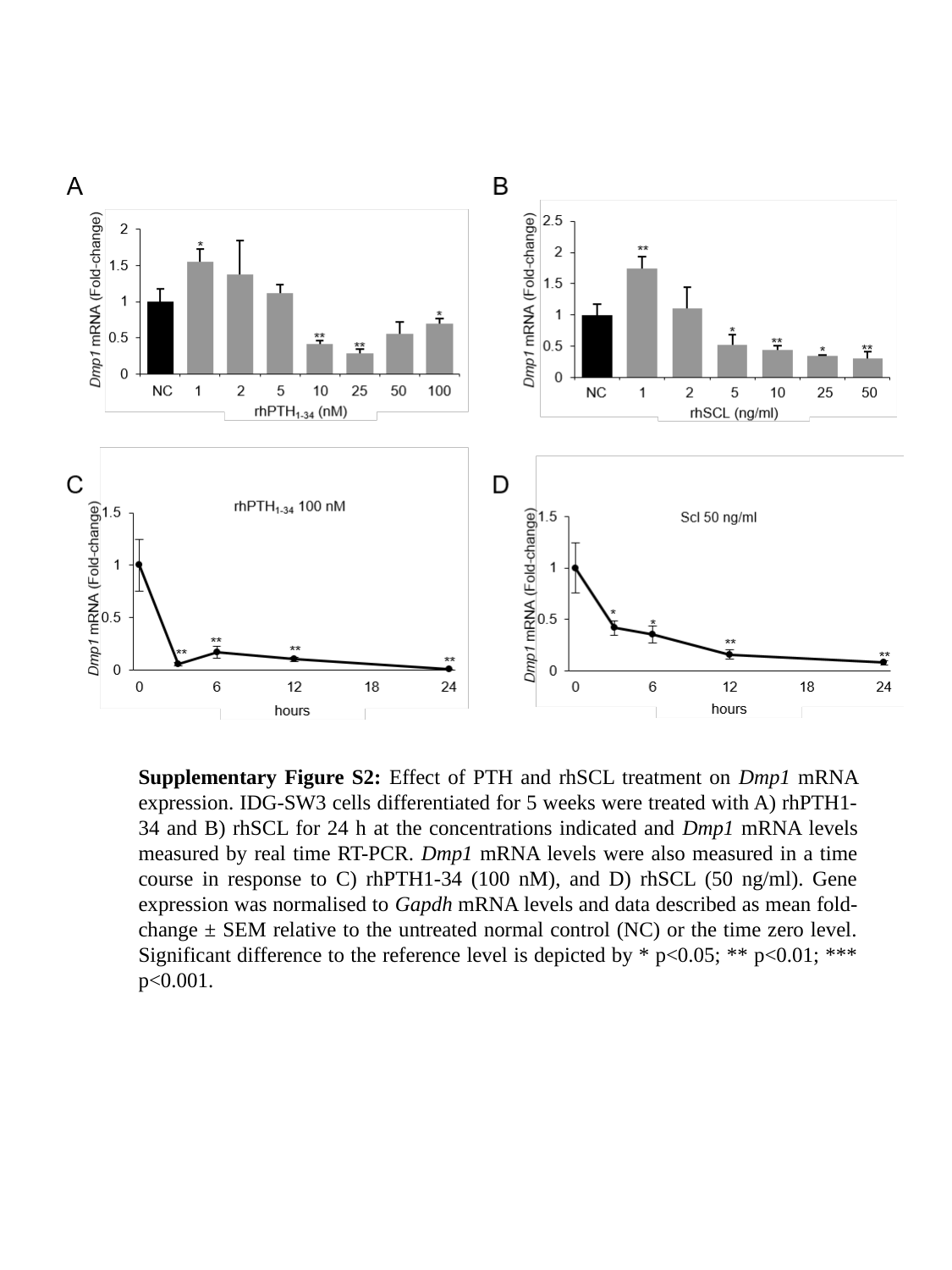

Supplementary Figure S2: Effect of PTH and rhSCL treatment on Dmp1 mRNA expression. IDG-SW3 cells differentiated for 5 weeks were treated with A) rhPTH1-34 and B) rhSCL for 24 h at the concentrations indicated and Dmp1 mRNA levels measured by real time RT-PCR. Dmp1 mRNA levels were also measured in a time course in response to C) rhPTH1-34 (100 nM), and D) rhSCL (50 ng/ml). Gene expression was normalised to Gapdh mRNA levels and data described as mean fold-change ± SEM relative to the untreated normal control (NC) or the time zero level. Significant difference to the reference level is depicted by * p<0.05; ** p<0.01; *** p<0.001.
